# A transcriptional complex of FtMYB102 and FtbHLH4 coordinately regulates the accumulation of rutin in *Fagopyrum tataricum*

**DOI:** 10.1101/2022.05.02.490379

**Authors:** Yaolei Mi, Yu Li, Guangtao Qian, Lucas Vanhaelewyn, Xiangxiao Meng, Tingxia Liu, Wei Yang, Yuhua Shi, Pengda Ma, Atia tul Wahab, Shilin Chen, Wei Sun, Dong Zhang

**Author notes:** These authors contributed equally to this work (Yaolei Mi, Yu Li).

## Abstract

Tartary buckwheat is rich in flavonoids, which not only play an important role in plant-environment interaction, but are also beneficial to human health. Rutin is a therapeutic flavonol which is massively accumulated in Tartary buckwheat. It has been demonstrated that transcription factors control rutin biosynthesis. However, the transcriptional regulatory network of rutin is not fully clear. In this study, through transcriptome and target metabolomics, we validated the role of FtMYB102 and FtbHLH4 TFs at the different developmental stages of Tartary buckwheat. The elevated accumulation of rutin in the sprout appears to be closely associated with the expression of FtMYB102 and FtHLH4. Yeast two-hybrid, transient luciferase activity and co-immunoprecipitation demonstrated that FtMYB102 and FtbHLH4 can interact and form a transcriptional complex. Moreover, yeast one-hybrid showed that both FtMYB102 and FtbHLH4 directly bind to the promoter of chalcone isomerase (*CHI*), and they can coordinately induce *CHI* expression as shown by transient luciferase activity assay. Finally, we transferred the FtMYB102 and FtbHLH4 into the hairy roots of Tartary buckwheat and found that they both can promote the accumulation of rutin. Our results indicate that FtMYB102 and FtbHLH4 can form a transcriptional complex by inducing *CHI* expression to coordinately promote the accumulation of rutin.

## INTRODUCTION

Flavonoids are major constituents of polyphenol in plant secondary metabolites. They consist mainly of anthocyanins, proanthocyanidins (PAs), flavonols, and flavones, which are present in almost all higher plants (Williams and Grayer, 2001). Flavonoids play critical roles in plant–environment interactions, including those involved in being uvioresistant, anti-herbivore and anti-pathogen (Emiliani *et al*. 2013; Barbehenn *et al*. 2011). Additionally, they also contribute to human nutrition and health (Williams *et al*. 2004). Rutin (quercetin-3-O-rutinoside), known as vitamin P, is a therapeutic flavonol bearing cytoprotective effects (Negahdari *et al*. 2021), which is massively accumulated in the seeds and leaves of Tartary buckwheat (Jiang *et al*. 2007). Previous studies have primarily focused on seeds, with few investigations on seedlings. The gene encoding flavonoid biosynthetic enzymes for the major flavonoid skeleton has been identified decades ago. Briefly, the pathway begins with phenylalanine and forms an important intermediate product, naringenin chalcone. Several key enzymes catalyze this precursor, such as phenylalanine aminolyase (PAL), cinnamic acid-4-hydroxylase (C4H), 4-coumarate-CoA ligase (4CL), and chalcone synthase (CHS) (Russell, 1971; Dixon & Lamb, 1979; Koes *et al*. 1989). Then, chalcone isomerase (CHI) catalyzes the resulting naringenin chalcone to form naringenin (Mona & Christopher, 1987). Furthermore, naringenin can be converted to flavones under the action of flavone synthase (FNS). Flavanone 3-hydroxylase (F3H), flavanone 3′-hydroxylase (F3′H), and flavonoid 3′5′-hydroxylase (F3′5′H) are responsible for producing dihydroflavonols which are precursors to flavonol and anthocyanidin branch (Hagmann *et al*. 1983; Cheng *et al*. 2014). The dihydroflavonols are subsequently catalyzed by flavonol synthase (FLS) to produce flavonols (Holton *et al*. 1993), among which rutin is a trademarked compound of Tartary buckwheat. *FtUGT73BE5* was characterized to be involved in rutin’s subsequent glycosylation (Yin *et al.,* 2020).

Additionally, the enzyme dihydroflavonol reductase (DFR), which is following anthocyanidin synthase (ANS) and anthocyanidin reductase (ANR), contribute to converting dihydroquercetin to proanthocyanidin (e.g., epicatechin) (Liew *et al*. 1998). Another pathway is the conversion of dihydroquercetin to catechin catalyzed by DFR and leucoanthocyanidin reductase (LAR).

Transcription factors (TFs) are the key factors to realize the spatiotemporal regulation patterns of secondary metabolites (Yang *et al*. 2012). Many TFs that regulate flavonoids synthesis have been reported, of which MYB is a crucial factor (Du *et al*. 2010; Vimolmangkang *et al*. 2013; Zhou and Memelink, 2016; Luo *et al*. 2018). MYB is one of the largest members of the plant transcription factor families and is classified into four types according to the number of MYB domain repeats (represented by R). The classes comprise of one MYB domain (R1/ R2-MYB), two (R2R3-MYB), three (R1R2R3-MYB), and four MYB domains (4R-MYB), among which R2R3 MYB is essential for flavonoids regulation (Dubos *et al*. 2010). Several MYB genes involved in flavonoids synthesis were first identified in *Arabidopsis thaliana*. For example, *PAP1/AtMYB75, AtMYB90, AtMYB113*, and *AtMYB114* in Arabidopsis from MYBA/SG6 play a positive regulatory role for anthocyanin accumulation by regulating *UFGT* and *DFR* expression (Zimmermann *et al*. 2004; Teng *et al*. 2005; Gonzalez *et al*. 2008; Lotkowska *et al*. 2015). The flavonol branch of the flavonoid biosynthesis is controlled in Arabidopsis by *MYB11, MYB12,* and *MYB111* from subgroup 7 of the R2R3-MYB family, which activates *CHS, CHI, F3H,* and *FLS*. However, *MYB3, MYB4, MYB7,* and *MYB32* with C2 repressors belonging to SG4 play an inhibitory role in the flavonoid synthesis pathway (Jin *et al*. 2000; Teng *et al*. 2005; Gonzalez *et al*. 2008; Zhou *et al*. 2015, 2017). In Tartary buckwheat, several MYBs have been reported to be involved in regulating the synthesis of flavonoids. Bai *et al*. (2014) reported that the overexpression of *FtMYB1* and *FtMYB2* significantly enhanced the accumulation of proanthocyanidins in *Nicotiana benthamiana* (*N. benthamiana*). *FtMYB31* is highly expressed in seeds and has been reported to positively regulate the biosynthesis of rutin in *N. benthamiana* (Hou *et al.,* 2021). Likewise, FtMYB11, and FtMYB13-16 act as negative regulators of rutin biosynthesis (Zhou *et al*. 2017; Zhang *et al*. 2018). Zhang *et al*. (2019) reported that light-induced *FtMYB116* can promote the accumulation of rutin. SG4-like *FtMYB18* and *FtMYB8* with tissue-specific expression patterns were recently identified as repressors of accumulating anthocyanin and proanthocyanidin (Huang *et al*., 2019; Dong *et al.,* 2020). However, the fine molecular mechanism of MYB’s regulation of flavonoids synthesis in Tartary buckwheat is still not well understood.

Generally, MYB recruits basic helix-loop-helix (bHLH) and WD40 proteins to form canonical MYB–bHLH–WDR (MBW) regulatory complexes to control flavonoid synthesis *in planta*. Several bHLHs modulating the anthocyanin and proanthocyanidin synthesis have been identified in dicots, including *AN1* and *JAF14* in Petunia (Spelt et al. 2000; Montefiori et al. 2015), *Delila* and *Mutabilis* in snapdragon (Shang et al. 2011), *TT8* and *GLABRA3* in *Arabidopsis* (Nesi et al. 2000; Payne et al. 2000), *MtTT8* in *Medicago truncatula* (Li et al. 2016). The pleiotropic bHLHs regulate the binding affinity of MYB to the *cis*-element of the target gene (Hichri et al., 2011) via interaction with the R3 repeat domain in R2R3 MYB proteins (Grotewold et al., 2000). Interestingly, MYB regulating flavonol synthesis requires bHLH in maize (Goff *et al.,* 1992), however, in Arabidopsis and grapevine it is independent of bHLH (Czemmel *et al*. 2009). However, relatively few bHLHs have been reported in Tartary buckwheat, with only FtTT8 reported to interact with other MYB TFs to regulate anthocyanin/proanthocyanidin synthesis (Huang *et al*. 2019; Dong *et al*. 2020; Wang *et al*. 2022). To date, the mechanism of bHLH regulation of flavonoids, as well as how MYB and bHLH synergistically regulate flavonoid synthesis remain largely elusive in Tartary buckwheat.

In this study, we identified and characterized two novel regulators, FtMYB102 and FtbHLH4, forming a transcriptional complex that co-regulates flavonoid synthesis, particularly flavonol, from sprout to seedling development. These results may have profound impacts on the understanding of precise transcriptional regulatory mechanisms of flavonoid production and even other secondary metabolites.

## MATERIALS AND METHODS

### Plant material and growth conditions

Two Tartary buckwheat varieties, TB115 and TB128, were used in this study. The growth conditions of Tartary buckwheat were previously reported (Zhang *et al.,* 2019). In brief, Tartary buckwheat seed coats were peeled off by soaking seeds in water for 20 min. Peeled seeds were then sterilized by washing with 75% ethanol for 1 min followed by 10 % sodium hypochlorite for 8 min and six 1-min washes with sterilized water. Next, the seeds were inoculated into sterile medium (Murashige and Skoog (MS) medium containing 0.8% agar and 1% sucrose). Subsequently, the sprouts and seedlings were harvested 3 and 8 days after sowing while kept at a photoperiod of 16h (light)–8h (dark) (~150μmolm^−2^ s^−1^) at 25°C, respectively. The plant samples were frozen immediately using liquid nitrogen and stored at −80°C for further use.

### RNA sequencing

The total RNA of Tartary buckwheat was extracted according to the manufacturer’s instructions (Tiangen, Beijing, China). RNA quality was then evaluated and the integrity number (RIN) of the samples was >8.0 using a Bioanalyzer 2100 instrument (Agilent Technologies, Palo Alto, CA, US). RNA sequencing was next conducted after establishing sequencing libraries using an IlluminaXten sequencing system (Illumina Inc., San Diego, CA, USA) as per the manufacturer’s instructions.

### RNA-seq data and phylogenetic analysis

After removing the low-quality bases and Illumina adapter sequences, approximately 46.9 G bases of clean data were utilized for this analysis. The STAR v.2.5 software was used to map these reads to the published Tartary buckwheat reference genome (Zhang *et al*. 2017a). Based on their FPKM (reads per kilobase of exon per million mapped reads) values (Mortazavi *et al*. 2008) differentially expressed genes (DEGs) were then sifted. FPKM values and DEG analysis (log_2_ Fold change ≥ 1) were analyzed using Htseq and DESeq2 software, respectively. The alignment of amino acid sequences with the default parameters and construction of phylogenetic tree was performed by using the neighbor-joining method via MEGA V.6 (Kumar *et al*. 2016). Bootstrap analyses with 1000 replications were used to assess the phylogenetic tree’s reliability nodes.

### Coexpression analysis

TFs were identified by using hmmscan software. Four genes, *C4H* (FtPinG0001575100), *F3H* (FtPinG0006662600), *F3′H* (FtPinG0002353900), and *CHI* (FtPinG0002790600), were used in coexpression analysis with a default value 0.05 to screen TFs that participated in the regulation of rutin synthesis. The paired genes were considered significantly coexpressed if the Pearson correlation coefficient (*r*) was greater than 0.95 (Ariani & Gepts, 2015).

### Quantitative RT-PCR

After the total RNA of Tartary buckwheat was isolated, the first-strand cDNA was synthesized using a commercial kit (TransGen, Beijing, CN) and then diluted to a concentration at 1 μg/ml for qRT-PCR using TransStart® Green qPCR SuperMix UDG (TransGen, Beijing, China) according to the manufacturer’s instruction. Additionally, qRT-PCR was conducted at Rotor-Gene Q (Qiagen, Hilden, Germany) . The experiment was conducted in triplicates, and the data were normalized to that of the reference gene, actin. The primers for qRT-PCR are listed in Table S1.

### Gene amplificaiton and vector construction

Using high fidelity DNA polymerase and cDNA as the template, transcriptional factors were amplified. Target gene promoters were amplified using the genome DNA of Tartary buckwheat as the template. The primers are listed in Table S1. These genes were constructed in a Blunt-Zero cloning vector (TransGen, Beijing, China) for further use.

### Yeast two-hybrid

The pLexA-FtbHLH4 reporter was cotransformed with the p8op-LacZ plasmid into the yeast strain EGY48, and positive clones were screened using the SD/-His/-Ura dropout media. After these clones were verified to be nonself-activated and non-toxic to yeast, they were mixed with the yeast strain, YM4271 transformed pB42AD-FtMYB102. Positive clones were obtained using SD/-His/-Ura/-Trp media with the addition of 20-mg/mL X-gal (5-Bromo-4-chloro-3-indolyl-β-d-galactopyranoside) for color development.

### Yeast one-hybrid

The AD-FtMYB102 effector and AD-FtbHLH4 were cotransformed with the *CHIp: LacZ* reporter into the yeast strain EGY48. After the transformants were screened for SD/-Trp-Ura dropout media, they were further inoculated on the dropout media containing X-gal for color variation.

### Co-immunoprecipitation

Briefly, bHLH4 and Myb102 fusions were transiently expressed in 5-week-old leaves of *Nicotiana benthamiana,* and infiltrated leaves were harvested 72 h after inoculation. Two grams of the corresponding leaves were ground in liquid nitrogen and inoculated in lysis buffer (50-mM Tris-MES, pH 8.0, 500-mM sucrose, 1-mM MgCl2, 10-mM EDTA, 5-mM dithiothreitol, 1-mM PMSF, and 1-mM Cocktail protease inhibitors) with end over end shaking for 30 min on ice. After centrifugation at 15,000 g for 1 h at 4°C, the supernatant was filtered and collected using MiraCloth. The protein concentrations of co-immunoprecipitation mixtures were then diluted to 1-mg/mL. Next, 500 μL of the above samples were precleared by incubating with protein G agarose beads at 4°C. After centrifugation, the precleared supernatant was incubated with protein G agarose beads conjugated rabbit-anti-GFP antibody (Yeason, CN) overnight at 4°C. Subsequently, the beads were collected by five times centrifugation-wash and eluted with 50-μL glycine aqueous solution (pH 2.5) for 1 min, and then 5-μL Tris pH 9.0 was added to neutralize. The eluted fractions and crude extracts were run in 10% SDS-PAGE gel and subjected to immunoblots with the corresponding antibodies. For the immunoblots, the primary monoclonal antibodies mouse-anti-Myc and mouse-anti-GFP (TransGen Biotech, CN) were used followed by secondary antibodies Goat-anti-Mouse-IgG horseradish peroxidase. An enhanced chemiluminescent reagent (StarLighter, CN) was used to detect the signal.

### Dual-luciferase reporter assay

The *35S-Ren-CHIp:LUC* reporter plasmid, construct *pGreen-62SK-FtMYB102,* and *pGreen-62SK-FtbHLH4* effector mixture were transformed into *Agrobacterium tumefaciens* strain EHA105, which were subsequently injected into the leaves of 5-week-old *N. benthamiana*. The tobacco leaves were harvested after growing under dark conditions overnight and subsequently under a 14 h light /10 h dark photoperiod for 4 days. The samples were handled as per instruction of the Dual-luciferase® Reporter Assay System (E1920, Promega, USA), and the signal was detected using SpectraMax i3x (Molecular Devices, USA).

### Luciferase complementation assay

The FtMYB102 and FtbHLH4 were constructed into plasmids of pCAMBIA1300-NLuc and pCAMBIA1300-CLuc, respectively. Both fusion proteins were then expressed in *N. benthamiana* by Agrobacterium-mediated transient expression. After the plants were placed in the dark for 1 day and photoperiod for 48 h, luciferase substrate and beetle luciferin (E1602, Promega, USA) at 150-μg/ml in saline were evenly sprayed on the back of *N. benthamiana* leaves. Luminescence was observed using NightSHADE LB985 (Berthold, Germany) after placing these leaves in the dark for 7 min.

### Transformation of the hairy roots of Tartary buckwheat

When the two pieces of cotyledon of Tartary buckwheat seedlings were unfolded, the hypocotyls and cotyledons, which were cut into ~0.5-cm segments and sheared into 0.5-cm pieces, respectively, were selected as explants. Then, these explants were infected with *Agrobacterium rhizogenes ACC10060* strain transformed target gene plasmids for 10 min after preculturing them on an MS solid medium for 1 day. Next, the explants were placed on an MS solid medium containing 500-mg/ml cefotaxime under 14 h light/10 h dark photoperiod at 25°C after coculturing them with bacteria for 3 days in the dark. The hairy roots were cut into 2–3 cm pieces and propagated in liquid MS medium (containing 30 g/L sucrose) at 25°C with a rotation speed of 80 rpm when they occurred and grew approximately 1–2 weeks later.

### Sample treatment and LC–MS analysis

Samples stored at −80°C were ground into fine powder. Then, 0.1 g of each sample was extracted in 1.0-mL 70% methanol aqueous at 4°C overnight and ultrasonicated for 30 mins. After centrifugation, 0.2 mL of each supernatant was filtered via a 0.22 μm membrane for liquid chromatography–mass spectrometry (LC–MS) analysis. The chromatographic conditions used for LC–MS were as the previous investigation (Yang, et al, 2020).

## RESULTS

### Flavonoid content varies in sprouts and seedlings

The secondary metabolites of plants have spatiotemporal characteristics, which means that the accumulation of secondary metabolites varies in different tissues and at different developmental stages. In addition to its seeds, Tartary buckwheat sprouts are also of great value. Therefore, we focused on the dynamic change in flavonoid content in Tartary buckwheat after germination. The sprouts and seedlings were harvested after sowing for 3 and 8 d, respectively. As shown in Figure 1A, the etiolated sprouts just emerged with their cotyledons still closed, yellow, and an apical hook. After 8 days, the cotyledons were fully open and green and the hypocotyls were fully grown. Target metabolomics showed more flavonoids, such as kaempferol, quercitrin, rutin, catechin, and epicatechin existed in the sprouts compared with the seedlings from two representative varieties TB115 and TB128 (Figure 1B & Table S2).

**Figure 1.**
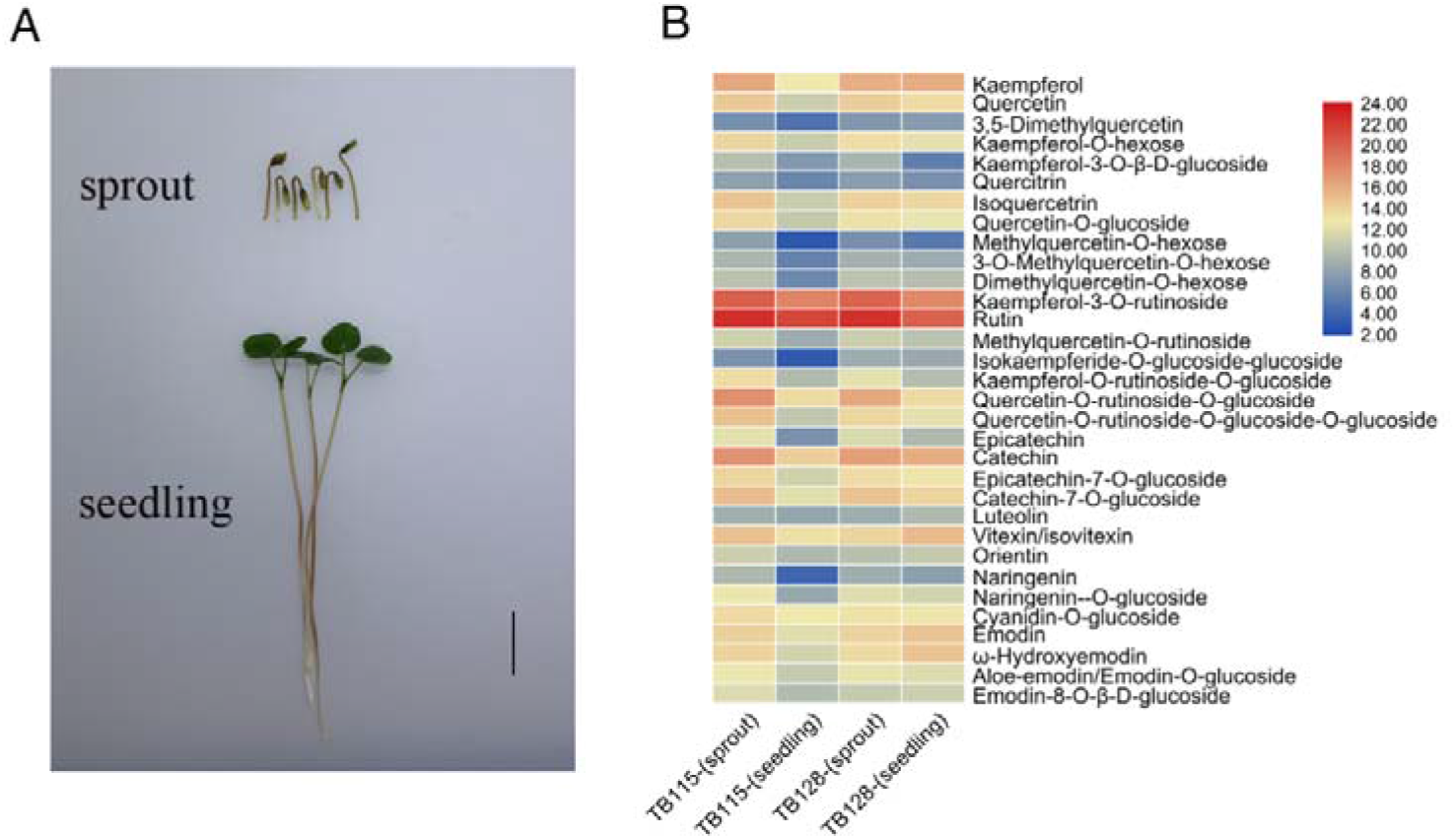
Flavonoid content in Tartary buckwheat at the sprout and seedling stages. (A) Photographs of Tartary buckwheat at the sprouting and seedling stages. Scale bar = 2□cm. (B) Heat map of the content of various flavonoids at the sprout and seedling stages of the TB115 and TB128 varieties. The depths of color in the red and blue rectangles indicate higher and lower flavonoid content level.

### Comparative analysis of transcriptional profiles

RNA sequencing was performed using the two varieties (TB115 and TB128) at the sprout and seedling stage to reveal the molecular mechanism that accounts for the differences in flavonoid contents of Tartary buckwheat at different developmental stages (accession number: PRJNA762576). We calculated the upregulated and downregulated genes of sprouts compared with seedlings. There were 3,215 and 3,084 genes upregulated in TB115 and TB128, respectively, with a total of 2,152 genes upregulated in both varieties (Figure 2A). The number of genes downregulated in TB115 and TB128 were 3,059 and 2,876, respectively, of which 1,855 genes were in common (Figure 2B). Following that, because flavonoid content was higher in sprouts than in seedlings, GO analysis was performed on the 2,152 upregulated genes (Figure 2C). The majority of the genes were discovered to be related to cell growth, which is consistent with the expectation that the seedlings were in the rapid growth stage. Interestingly, we discovered that the phenylpropanoid and secondary metabolic pathways both had a substantial number of annotated genes, which confirmed our previous metabolome results, indicating that a large number of secondary metabolites change in plants during these two stages.

**Figure 2.**
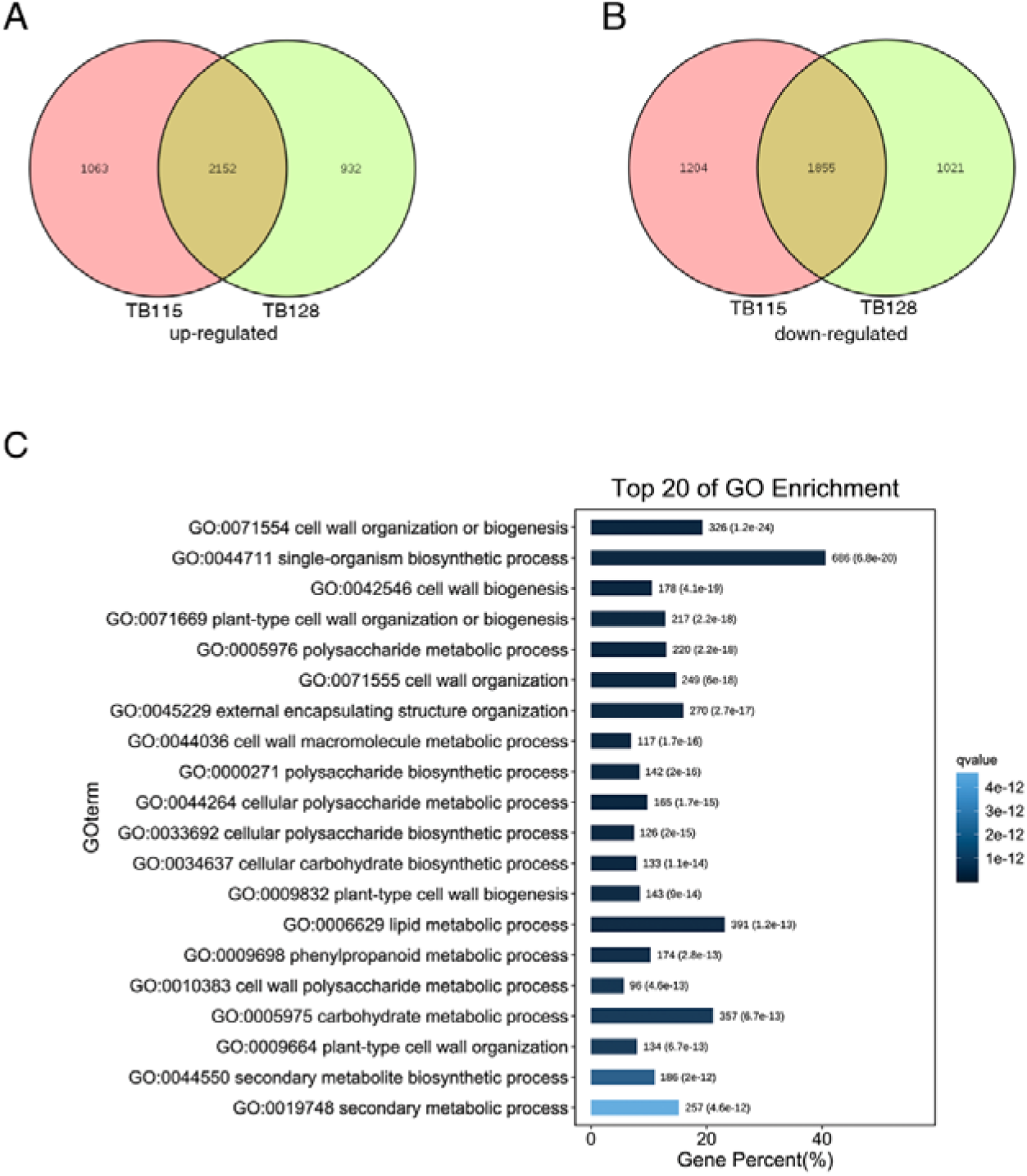
Gene expression varied in sprouts from seedlings in two varieties TB115 and TB128, and GO analysis among these differential genes. (A) Upregulated genes in the two varieties. Among these genes, 2,152 genes (brown) were upregulated in both varieties. (B) 3,059 and 2,876 genes were downregulated genes in both varieties. 1,855 genes (brown) were overlapped. (C) GO analysis on the 2,152 upregulated genes. Top 20 of the GO enrichment were listed.

### Expression of genes related to flavonoid biosynthesis

We specifically analyzed the expression of genes related to flavonoid synthesis. The results showed that the expression of most of these genes in sprouts was higher in sprouts than in seedlings (Figure 3 & Table S3). For example, two out of the five transcripts of *PAL* (FtPinG0001546000 and FtPinG0005713400), three *CHS* (FtPinG0000551600, FtPinG0000551900, and FtPinG0008806400) and two *4CL* (FtPinG0003957300 and FtPinG0005072700) displayed a much higher expression level in the sprouts compared to seedlings. Additionally, *C4H* (FtPinG0001575100), *F3′H* (FtPinG0002353900), *F3H* (FtPinG0006662600 and FtPinG0008251700), *DFR* (FtPinG0002371500), *FLS* (FtPinG0006907100), *CHI* (FtPinG0002790600), and *GTR* (FtPinG0006606900) displayed a similar pattern. The expression patterns of these flavonoid structure genes were consistent with the flavonoid content described previously (Figure 1B), indicating that sprouts accumulate more flavonoids than seedlings. After that, we performed real-time quantitative RT-PCR for further validation of the expression of flavonoid structure genes. The results showed that the expression levels of these genes in the sprouts were significantly higher than those in seedlings (Figure S1), which was also consistent with our transcriptome results.

**Figure 3.**
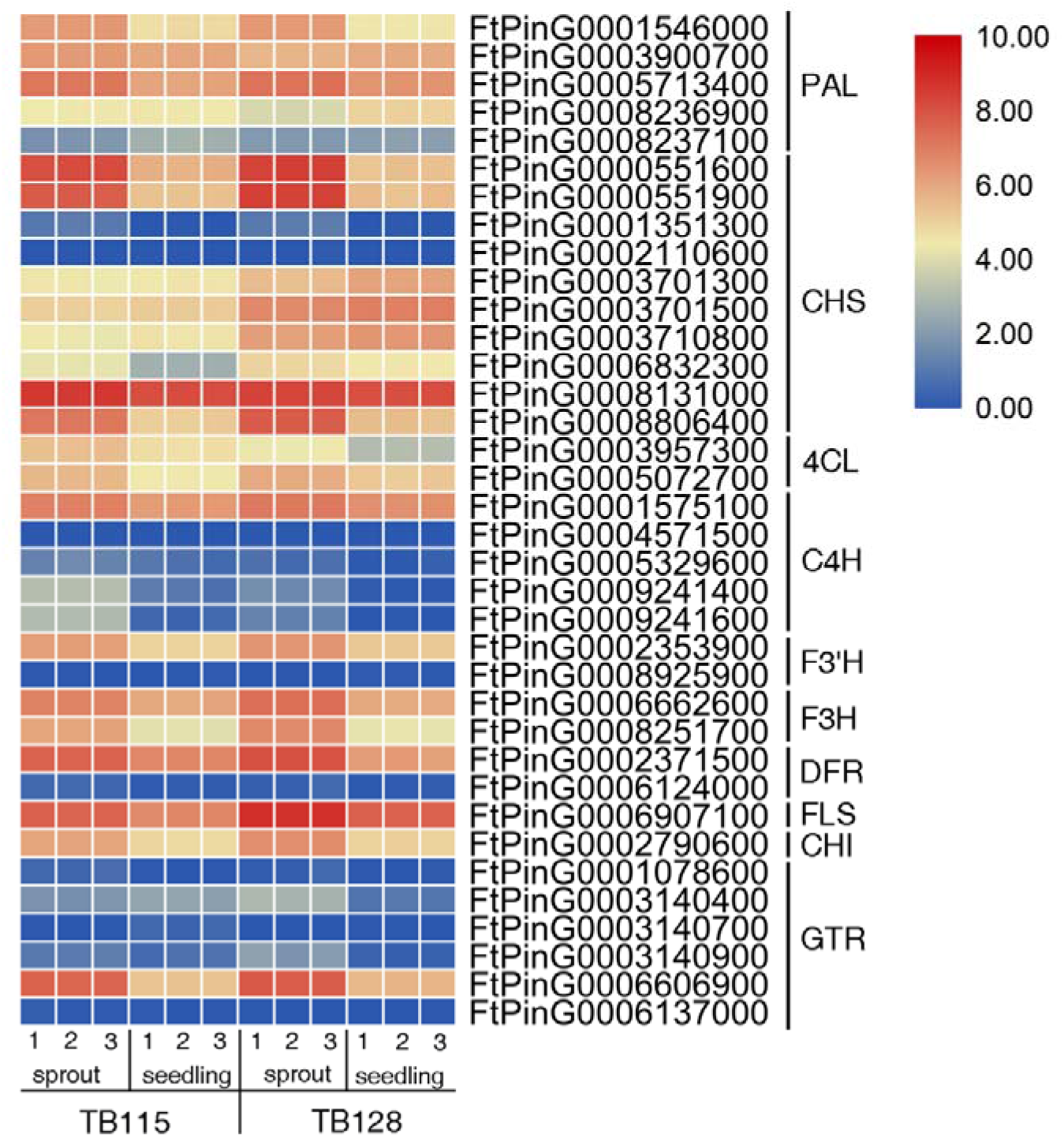
Heat map of the expression levels of flavonoid synthesis genes at sprout and seedling stages. For each variety, the sprouting and seedling level were studied, the numbers (1-3) represent three biological repeats. The depths of color in the red and blue rectangles indicate higher and lower z scores of RNA expression level. PAL: phenylalanine ammonialyase; CHS: chalcone synthase; 4CL: 4-coumarate:CoA ligase; C4H: cinnamate-4-hydroxylase; F3′ H: flavonoid-3′-hydroxylase; F3H: flavanone-3-hydroxylase; DFR: dihydroflavonol reductase; FLS: flavonol synthase; CHI: chalcone isomerase; GTR: glucosyl transferase

### Coexpression revealed that FtMYB102 directly regulated *CHI* expression

To obtain the potential TFs involved in flavonoid biosynthesis, coexpression was performed to identify the TFs, which are tightly coexpressed with four highly expressed structural genes: *C4H* (FtPinG0001575100), *F3H* (FtPinG0006662600), *F3′H* (FtPinG0002353900), and *CHI* (FtPinG0002790600). We identified 19, 29, 29, and 41 TFs that were coexpressed among *C4H, CHI, F3H*, and *F3′H*, respectively, through this analysis (Table S4 and S5). Among these TFs, 14 were tightly coexpressed with all the four genes (Figure 4A). Similarly, all these 14 TFs exhibited higher expression in sprouts than in seedlings in a pattern similar to *C4H, CHI, F3H*, and *F3′H* (Figure 4B), where the largest gene family (4 MYBs out of 14 TFs) were proposed to directly regulate the synthesis of flavonoids.

**Figure 4.**
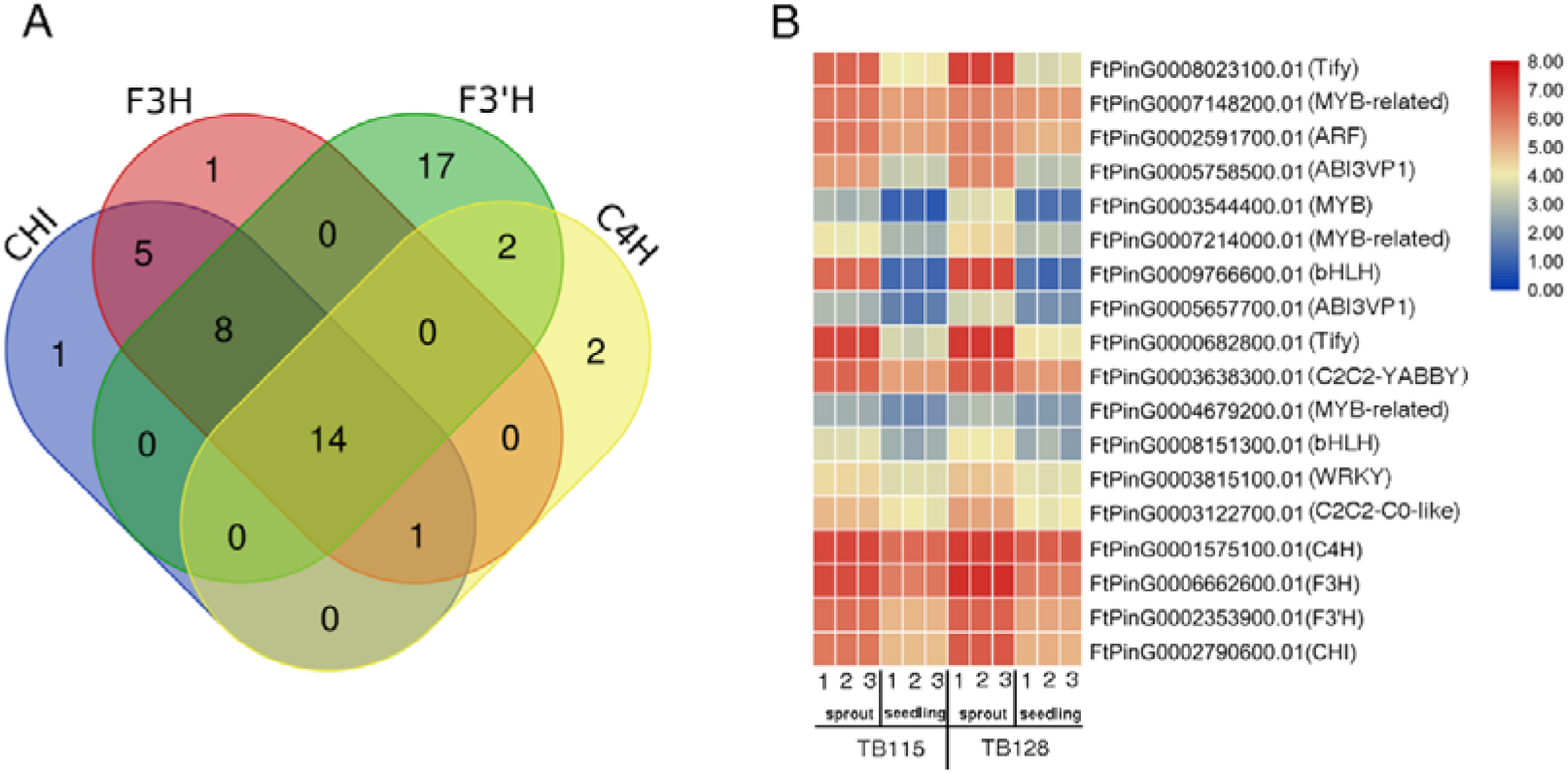
Identification of transcription factors coexpressed with flavonoid synthesis genes. (A) The Venn diagram shows the number of genes coexpressed with CHI, F3H, F3′H, and C4H. (B) Heat map of the expression levels of the 14 TFs indicated between brackets, including CHI, F3H, F3′H, and C4H. The numbers one, two, and three represent three biological repeats. The depths of color in the red and blue rectangles indicate higher and lower z scores of RNA expression level.

MYB TFs that regulated flavonoid synthesis were also identified using the yeast one-hybrid assay (Y1H). The promoters of *C4H, CHI, F3H,* and *F3′H* were cloned into pLacZ-2μ reporter vectors in fragments (2,000 bp), while the four MYB TFs were cloned into pB42AD (GAL4 activation domain [AD] fused). FtPinG0007148200.01, which we termed FtMYB102, is bound to the promoter of *CHI,* but not to the promoters of the other three examined genes, as shown in Figure 5A (data not shown). Sprouts accumulated more *FtMYB102* transcripts than seedlings (Figure 5B). Further analysis showed FtMYB102 coded for 265 amino acids and the phylogenetic relationship revealed that FtMYB102 clustered with AtMYB5 (Figure 6A & Figure S2A), regulating the accumulation of PA (Deluc *et al*. 2008) together with *PhPH4*(*Petunia hybrida*), MdMYB12(*Malus domestica*), and VvMYB5b(*Vitis vinifera*), which belonged to classic R2R3 MYB TFs (Figure 6B).

**Figure 5.**
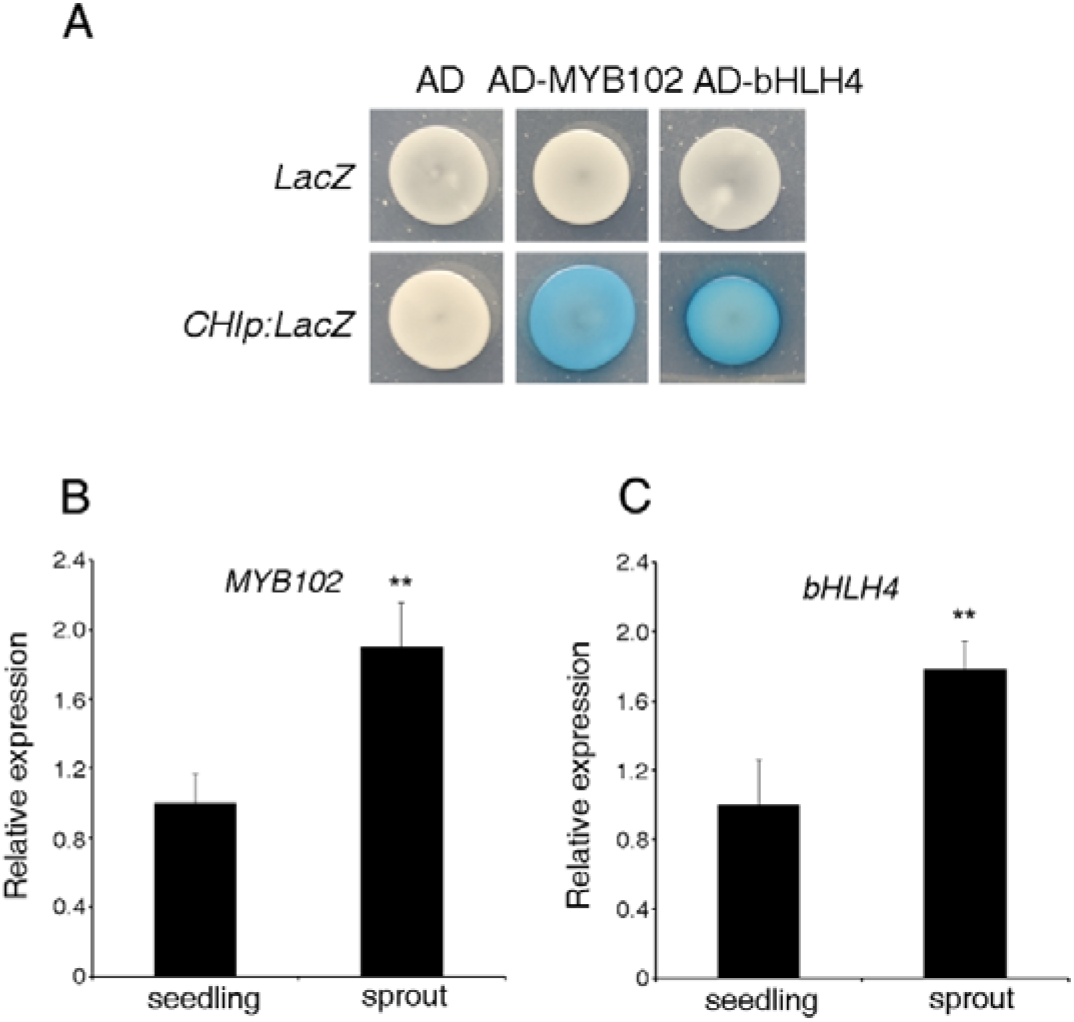
FtMYB102 and FtbHLH4 are candidate transcription factors that regulate rutin synthesis. (A) Yeast one-hybrid assay showing that AD-MYB102 and AD-bHLH4 bind to the promoter regions of *CHI*. (B) and (C) MYB102 and *bHLH4* expression in sprout and seedling. The relative expression levels were normalized to those of the actin control. Data represent the means ± standard deviation of biological triplicates.

**Figure 6.**
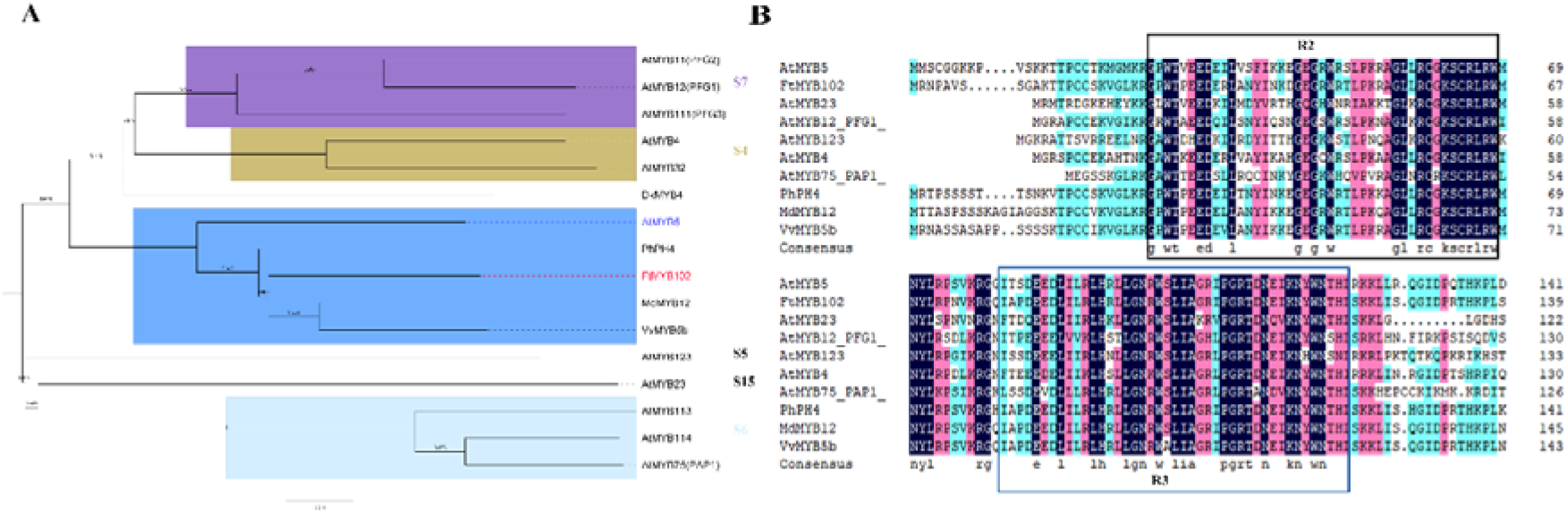
Phylogenetic and structural domain analysis of FtMYB102. (A) Phylogenetic analysis of FtMYB102 with MYB TFs in different subgroups and species. Three S7 group members: AtMYB11-12 and AtMYB111 in *A. thaliana;* two S4 group members: AtMYB4 and At MYB32; one S5 and S15 group member AtMYB123 and AtMYB23, respectively; three S6 group members AtMYB75 and AtMYB113-114. FtMYB102, PhPH4(*Petunia hybrida*), MdMYB12(*Malus domestica*) and VvMYB5b(*Vitis vinifera*) were classified as AtMYB5. (B) The structural domain analysis of FtMYB102 showed it was a classic R2R3 MYB TF.

### bHLH candidate genes involved in flavonoid synthesis

Generally, bHLH TFs are able to interact with MYB TFs to form a transcription complex that regulates the expression of flavonoid synthesis genes. It was previously reported in *Arabidopsis thaliana* that TT8 can form a complex with the MYB transcription factor to regulate the synthesis of flavonoids (Zhou *et al*. 2012). In this study, we identified seven homologous genes named *FtbHLH1–FtbHLH7* in Tartary buckwheat based on the TT8 sequence alignment in *A. thaliana*. Then, we performed a phylogenetic analysis of these seven genes to construct an evolutionary tree with other Arabidopsis bHLH TFs (Figure S2B). Among them, FtbHLH1 and FtbHLH3–4 were classified as IIIf, while FtbHLH2 and FtbHLH5–7 were classified as III(d+e). Interestingly, many bHLH genes reported in *A. thaliana* and rice that were involved in flavonoid synthesis belonged to the IIIf subfamily, (Ludwig *et al*. 1989; Feyissa *et al*. 2009; Zhou *et al*. 2012), which indicated that FtbHLH1, FtbHLH3-4 were involved in the synthesis of flavonoids.

To further verify whether the three bHLH TFs participated in the regulation of flavonoids, Y1H was used to clarify whether they can directly bind to the promoters of the flavonoid synthesis genes. As described above, fragments (2,000 bp) of the promoters of *C4H, CHI, F3H,* and *F3′H* were cloned into pLacZ-2μ reporter vectors, whereas the three bHLH TFs were cloned into pB42AD. FtbHLH4 bound to the *CHI* promoter but not to the other three promoters as shown in Figure 5A, whereas FtbHLH1 and FtbHLH3 could not bind to any promoters (data not shown). As a result, we focused our research on FtbHLH4. Similar to FtMYB102, we found that the expression of FtbHLH4 was significantly higher in sprouts than in seedlings (Figure 5C). As shown, the expression patterns of FtMYB102 and FtbHLH4 genes were found to be consistent with those of flavonoid synthesis genes, suggesting that the two genes directly regulated the expression of flavonoid synthesis genes.

### FtbHLH4 physically interacts with FtMYB102

To identify whether MYB102 and bHLH4 can form a complex, yeast two-hybrid experiment (Y2H) was performed by fusing FtbHLH4 with the LexA DNA-binding domain and the FtMYB102 with the B42 activation domain (AD), respectively. Y2H results showed that LacZ reporter expression was strongly induced when both FtbHLH4 and FtMYB102 were simultaneously transformed into yeast, but either individual or none was effective, indicating FtbHLH4 could interact with FtMYB102 in yeast (Figure 7A). Next, a transient luciferase activity assay was performed in *N. benthamiana* to further examine the interaction between FtbHLH4 and FtMYB102. As shown in Figure 7B, no or weak luciferase fluorescence signal was observed in leaves of *N. benthamiana* injected with water, nLUC+cLUC, nLUC+ FtbHLH4-cLUC, and FtMYB102-nLUC+cLUC. However, when leaves were injected with FtMYB102-nLUC+ FtbHLH4-cLUC, there was a significant increase in the fluorescence value of leaves.

**Figure 7.**
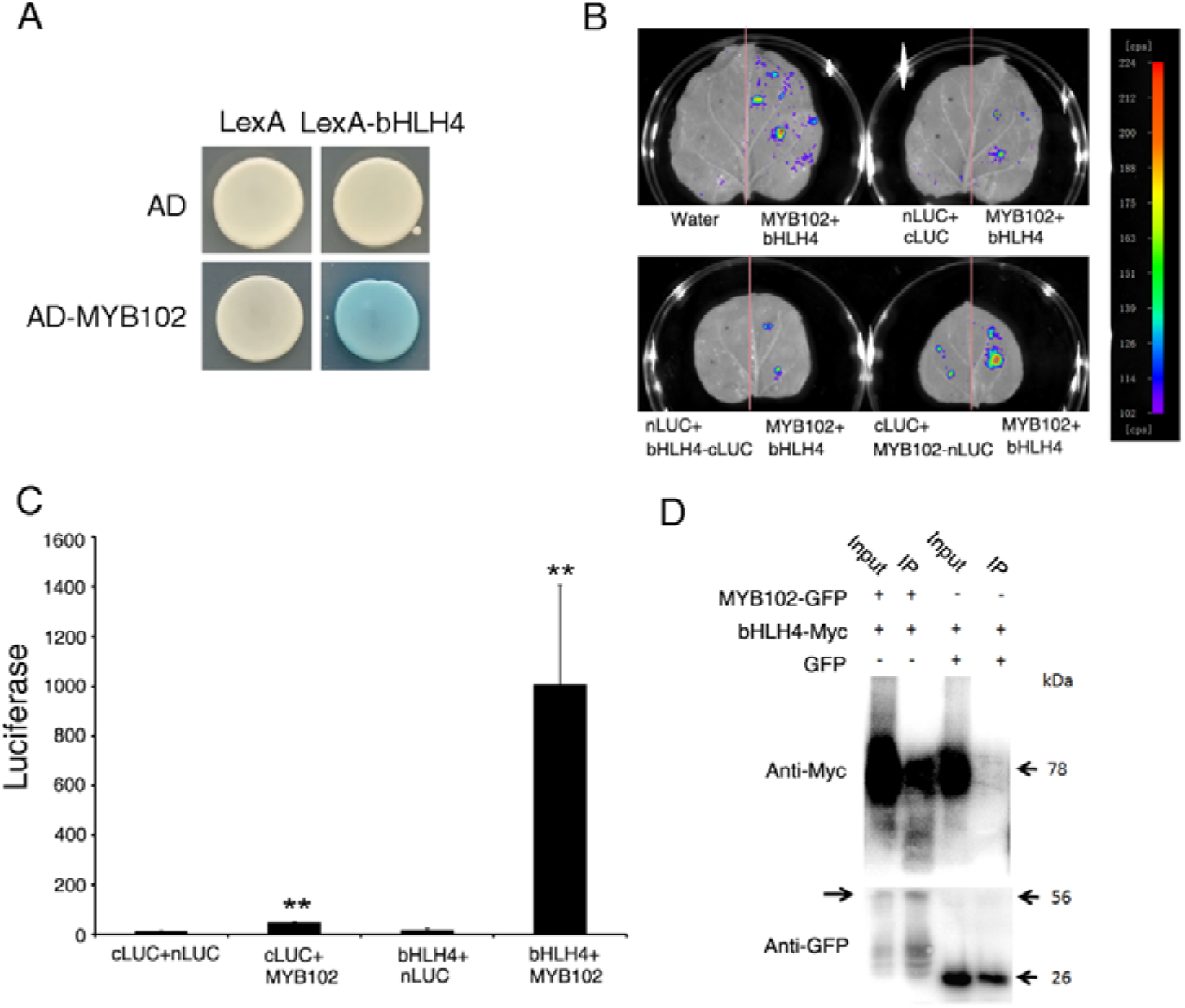
FtbHLH4 physically interacts with FtMYB102. (A) Yeast two-hybrid assay using bHLH4 and MYB102 constructs with AD: the B42 activation domain alone; AD-MYB102: MYB102 fused with the B42 activation domain; LexA: the LexA DNA-binding domain alone; LexA-bHLH4: the LexA DNA-binding domain fused to bHLH4. (B) Luciferase Complementation Imaging Assay (LCI assay) of MYB102–nLUC with bHLH4-cLUC in tobacco leaves with Water, nLUC+ cLUC, nLUC+ bHLH4-cLUC and cLUC+ MYB102–nLUC served as controls; MYB102–nLUC + bHLH4-cLUC in the four leaves were four biological repeats. (C) The interaction between MYB102 and bHLH4 was quantitatively detected using LCI. Data represent the means ± standard deviation of biological triplicates. (D) Co-IP assay with MYB102-GFP and bHLH4-Myc coexpression in tobacco leaves. +: the corresponding component was added to the reaction system; –: no corresponding component added; the number on the right represents the size of the proteins.

The signal of FtMYB102-nLUC+ FtbHLH4-cLUC increased more than 400 times when compared to the control groups (Figure 7C). Furthermore, we conducted co-immunoprecipitation assays to verify the FtbHLH4–FtMYB102 interaction. To achieve this assay, 35S:FtMYB102-MYC was coinfiltrated with 35S:GFP and 35S:FtbHLH4-GFP inserted into *N. benthamiana* leaves. Afterward, the total plant proteins were extracted. Immunoprecipitation was conducted with the GFP antibody linked to agarose beads, where the FtMYB102-MYC fusion protein was pulled down in samples coexpressing FtbHLH4-GFP, but not GFP alone (Figure 7D). Therefore, these results indicate that FtbHLH4 interacted with FtMYB102.

### FtMYB102 and FtbHLH4 Coordinate Activation the Expression of *CHI*

Previous studies have suggested that MYB and bHLH formed complexes that regulated the expression of flavonoids synthesis genes. To study the regulation pattern of downstream gene expression by FtMYB102 and FtbHLH4, a dual-luciferase reporter assay was performed in *N. benthamiana* (Figure 8A). Figure 8B showed that there was no LUC signal difference between 62SK (*Agrobacterium tumefaciens* strain GV3101 harboring recombinant plasmids), FtMYB102-62SK, FtbHLH4-62SK, and FtMYB102-62SK+ FtbHLH4-62SK co-transformed with empty *pGreenII 0800-LUC* vector into leaves, respectively. However, when co-transformed with *ProFtCHI*:*LUC*, the LUC signal in the individual transformation with FtMYB102-62SK and FtbHLH4-62SK was increased significantly than the 62SK control. In addition, the LUC signal in the co-transformation of FtMYB102-62SK and FtbHLH4-62SK was further increased. Therefore, the results indicated that FtbHLH4 and FtMYB102 can individually and coordinately activate *CHI* expression via a complex.

**Figure 8.**
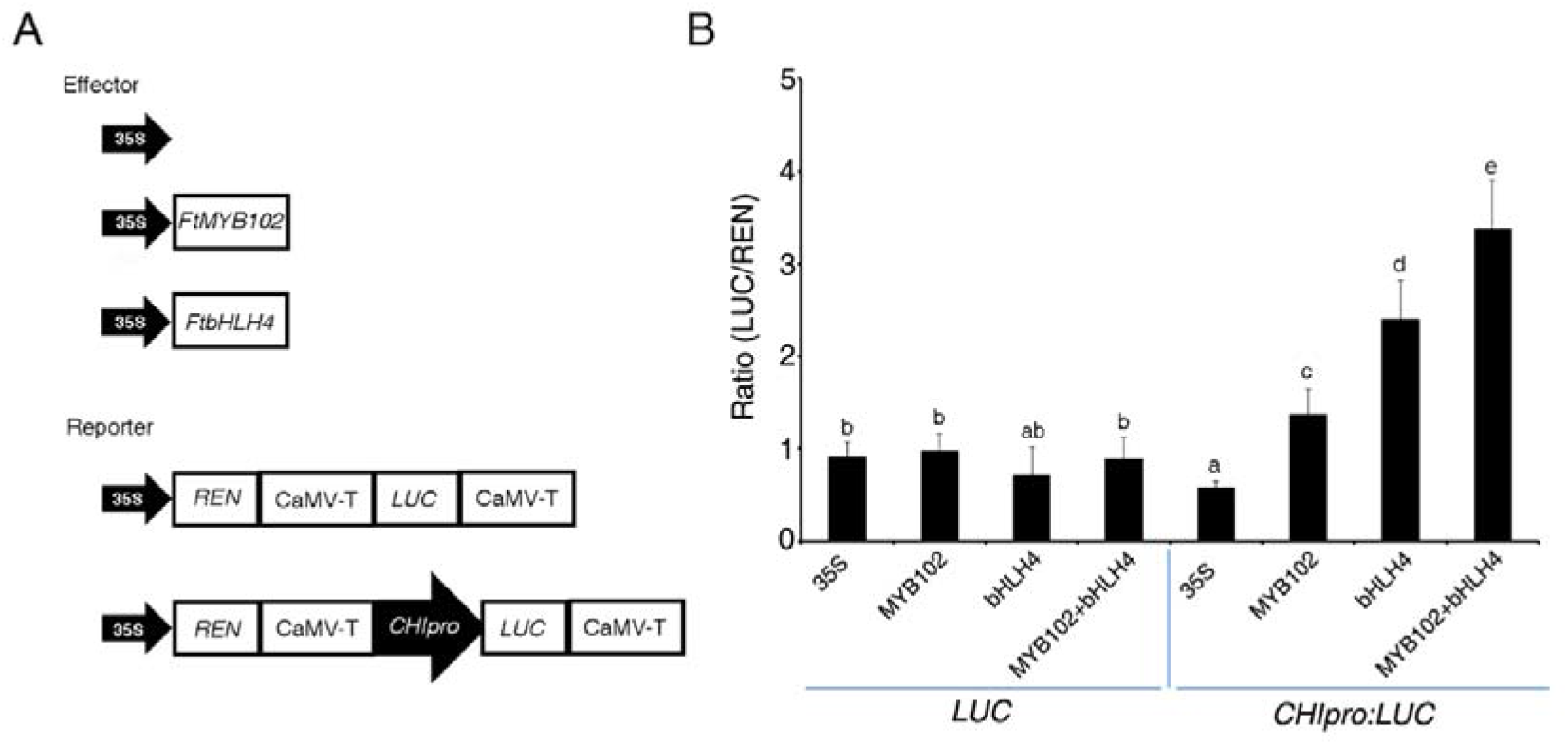
MYB102 and bHLH4 coordinately activate the expression of *CHI*. (A) Schematic diagrams of the effector and reporter plasmids used in dual-LUC assays. REN, Renilla luciferase; LUC, firefly luciferase. (B) Dual-LUC assay in tobacco leaves using the constructs shown in (A). The 35S effector was used as a negative control. Data represent the means ± standard deviation of biological triplicates.

### Overexpression of both FtMYB102 and FtbHLH4 can promote flavonoids biosynthesis

To verify whether FtMYB102 and FtbHLH4 promoted the synthesis of flavonoids, we generated transgenic Tartary buckwheat hair roots that overexpressed *FtMYB102* (OE-FtMYB102) and *FtbHLH4* (OE-FtbHLH4). Three independent transgenic hairy root lines were used for further functional analysis. The expression of *FtMYB102* or *FtbHLH4* in the three lines was significantly higher than that in the WT, showing that both the genes were overexpressed in the transgenic hair roots according to qRT-PCR results (Figure S3). Concomitantly, LC–MS analysis showed OE-FtMYB102 and OE-FtbHLH4 produced more flavonoids than WT (except catechin in MYB102OX-4 and MYB102-OX-6) (Figure 9A). The overexpressing lines displayed a significant increase in rutin, indicating that FtMYB102 and FtbHLH4 had greatly promoted rutin synthesis. The qRT-PCR analysis consistently revealed enhanced expressions of *CHS, CHI, F3H,* and *FLS* in OE-FtMYB102 and OE-FtbHLH4 lines compared with the control (Figure 9B), suggesting that FtMYB102 and FtbHLH4 promote flavonoid synthesis through these four genes.

**Figure 9.**
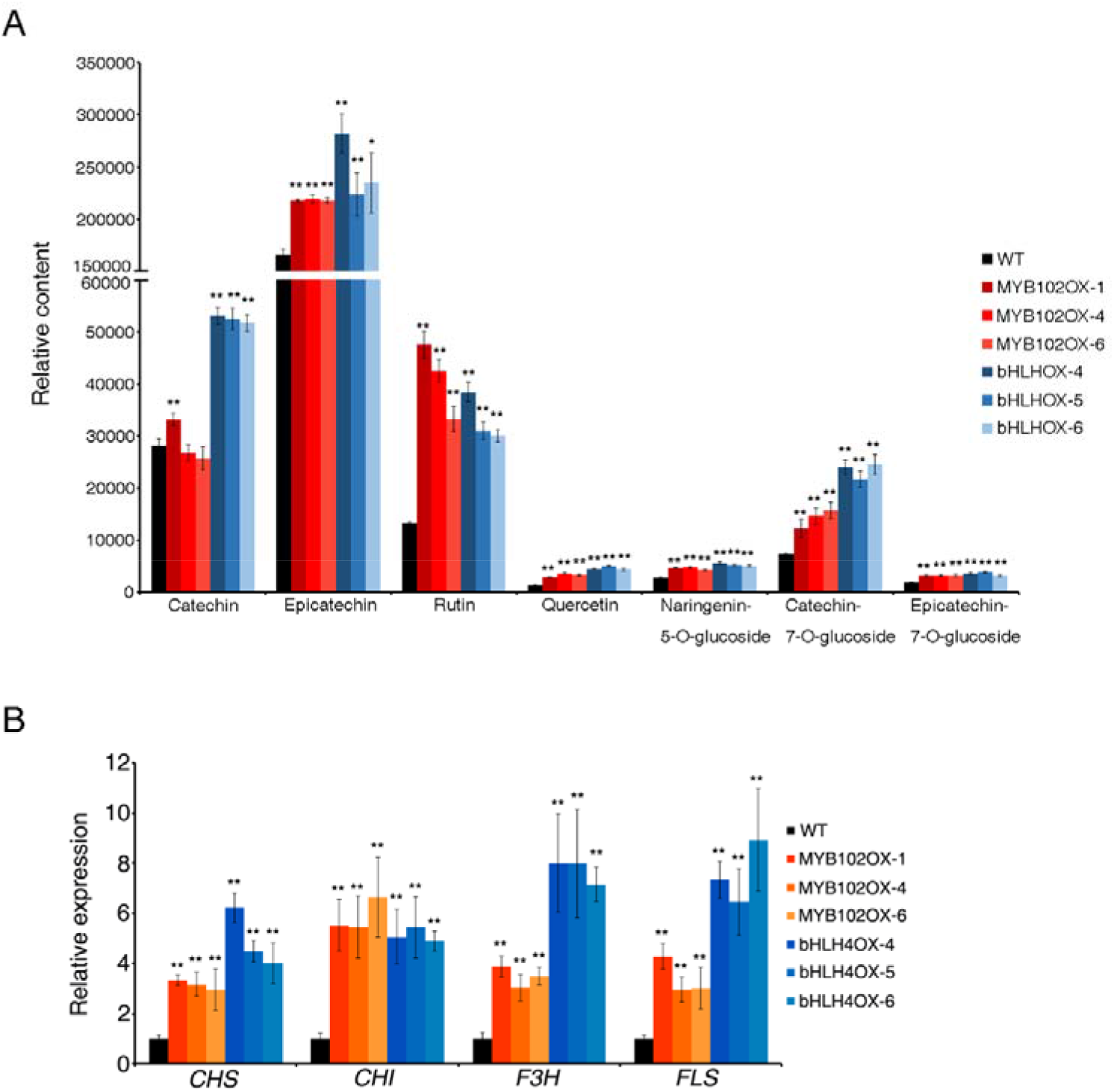
Overexpression of MYB102 and bHLH4 can promote flavonoid biosynthesis in Tartary buckwheat hairy roots. (A) Relative flavonoid content in MYB102 and bHLH4 overexpression lines in Tartary buckwheat hairy roots. Double asterisks indicate a significant difference at P < 0.01 using the Student’s t test-test. (B) Detection of gene expression related to flavonoid synthesis pathways in MYB102 and bHLH4 transgenic lines. The relative expression levels were normalized to that of the actin control. Data represent the means ± standard deviation of biological triplicates.

## DISCUSSION

Tartary buckwheat is rich in flavonoids, especially its trademark rutin, and has a high nutritional and medicinal value. This study showed that in addition to Tartary buckwheat seeds which are commonly consumed (Yang *et al*. 2020), the flavonoid content of its sprouts and seedlings is considerably high. Given that this vegetable has been gaining popularity in China, understanding its edible and medicinal value is important. Secondary metabolite biosynthesis in plants is not only tissue-specific but also dependent on the plant’s developmental stages. According to Czechowski *et al*. (2016), the content of artemisinin increased gradually during the development of a young leaf to its mature form. Arce-Rodríguez *et al*. (2017) showed that capsaicinoid accumulation is dependent on the developmental stage of chili pepper fruits. As a result, two Tartary buckwheat varieties were used in this study to investigate the flavonoid content during two developmental stages (sprouts and seedlings). The results of the metabolome showed that the content of most flavonoids was higher in the sprouts than in the seedlings. Most previous studies on the flavonoid content of Tartary buckwheat have focused on seeds. As a result, this study is critical in determining the flavonoid content of Tartary buckwheat in the different developmental stages. Follow-up consumption studies might be an interesting research topic.

To reveal the transcriptional mechanism of flavonoid accumulation at the sprout and seedling stages, RNA sequencing and coexpression analysis was performed. TFs coexpressed with four flavonoid synthetic genes (*CHI, F3H, F3′ H*, and *C4H*) were screened. The observations showed that one of the MYB TFs, namely FtMYB102, showed a consistent expression pattern with all the four genes. Y1H showed that FtMYB102 directly binds to the promoter of *CHI*. Then, a transient luciferase activity assay was conducted in *N. benthamiana* to demonstrate that FtMYB102 promoted *CHI* expression (Figure 8B). A new MYB transcription factor was discovered here that regulated flavonoid synthesis at different Tartary buckwheat development stages. FtMYB102 clustered with AtMYB5, which does not belong to the SG7 group of MYBs that particularly control flavonol accumulation, in an unusual phylogenetic relationship (Wang *et al*. 2017). AtMYB5 and its homology in grape, VvMYB5b, have generally been considered a regulator of PA (Deluc *et al*. 2008). The overexpression of FtMYB102 not only significantly promoted PA accumulation but also flavonol accumulation in this study (Figure 9A), which further broaden the roles of MYB5 genes on flavonoid synthesis regulation.

Generally, bHLH proteins act as transcriptional factors, subtly controlling flavonol metabolism via the affinity of *cis*-regulatory element of downstream genes (Li *et al*. 2020), broad or restricted expression pattern, influenced by their dimerization properties with MYB TFs (Feller *et al*. 2011). They directly bind to the *cis*-elements (G-box and E-box) of structural genes via the basic region, located at the N-terminal end of the domain, while the HLH region, at the C-terminal end, is involved in homo- and hetero-dimerization (Toledo-Ortiz et al. 2003). In this study, we detected seven homologs of AtTT8 in the Tartary buckwheat genome, which exert distinct DNA-binding functions, i.e., only FtbHLH4 binds to *CHI* promoter but others are not able to, which could explain the specific binding of bHLH members to *cis*-elements to regulate flavonoid synthesis. FtbHLH4 was found to be more highly expressed in sprouts than in seedlings, and it resembled FtMYB102, suggesting that FtbHLH4 regulates sprout and seedling flavonoid biosynthesis. The regulation of flavonoids in Tartary buckwheat was also confirmed by the overexpression of *FtbHLH4* in hairy roots, which leads to the accumulation of flavonoids.

It has also been reported that MYB TFs can interact with bHLH to form a complex that regulates gene expression in *Arabidopsis thaliana* (Gonzalez *et al*. 2008); however, this has not been reported in Tartary buckwheat. In this study, Y2H, transient luciferase activity assay, and CoIP demonstrated that FtbHLH4 can interact with the FtMYB102 (Figure 7). Furthermore, through transient luciferase activity assay, it was shown that when both MYB102 and bHLH4 TFs were present, the activation of *CHI* was significantly stronger than when they were present independent of each other (Figure 8). These results indicate that MYB102 and bHLH4 interacted with each other to form a complex that directly bound to the *CHI* promoter and actively triggered *CHI* gene expression. Our results strongly suggest that the combined action of FtMYB102 and FtHLH4, which enhances target gene *CHI* expression, is critical for high flavonoid production (Figures1, 7–9). These findings highlight the importance of the interaction specificity between the cooperative partners of FtMYB102 and FtHLH4 proteins in regulating flavonoid metabolism at the sprout and seedling stages.

Based on the results, we developed a working model (Figure 10). In the sprout stage, the highly expressed MYB102 and bHLH4 interact to form a transcriptional complex that directly binds to the promoter of *CHI* and induces its expression. *CHI* is an essential catalytic enzyme in the rutin synthesis pathway. Under its catalysis, naringenin chalcone is transformed into chalcone, and rutin is further synthesized under the action of several other catalytic enzymes. Therefore, when the expression level of *CHI* is high, the synthesis of rutin is increased, resulting in the formation of high rutin in sprouts. In the seedling stage, the expression of MYB102 and bHLH4 decreased, resulting in decreased *CHI* expression, ultimately reducing the content of rutin in the seedlings. It remains unknown what factors cause the difference in the transcription levels of MYB102 and bHLH4 when Tartary buckwheat is in different developmental stages. Since many differences are observed when plants are at different stages of development, the changes in the expression of these two TFs could have been produced by a hormone or a single signal molecule. Hence, further research is needed to address this issue.

**Figure 10.**
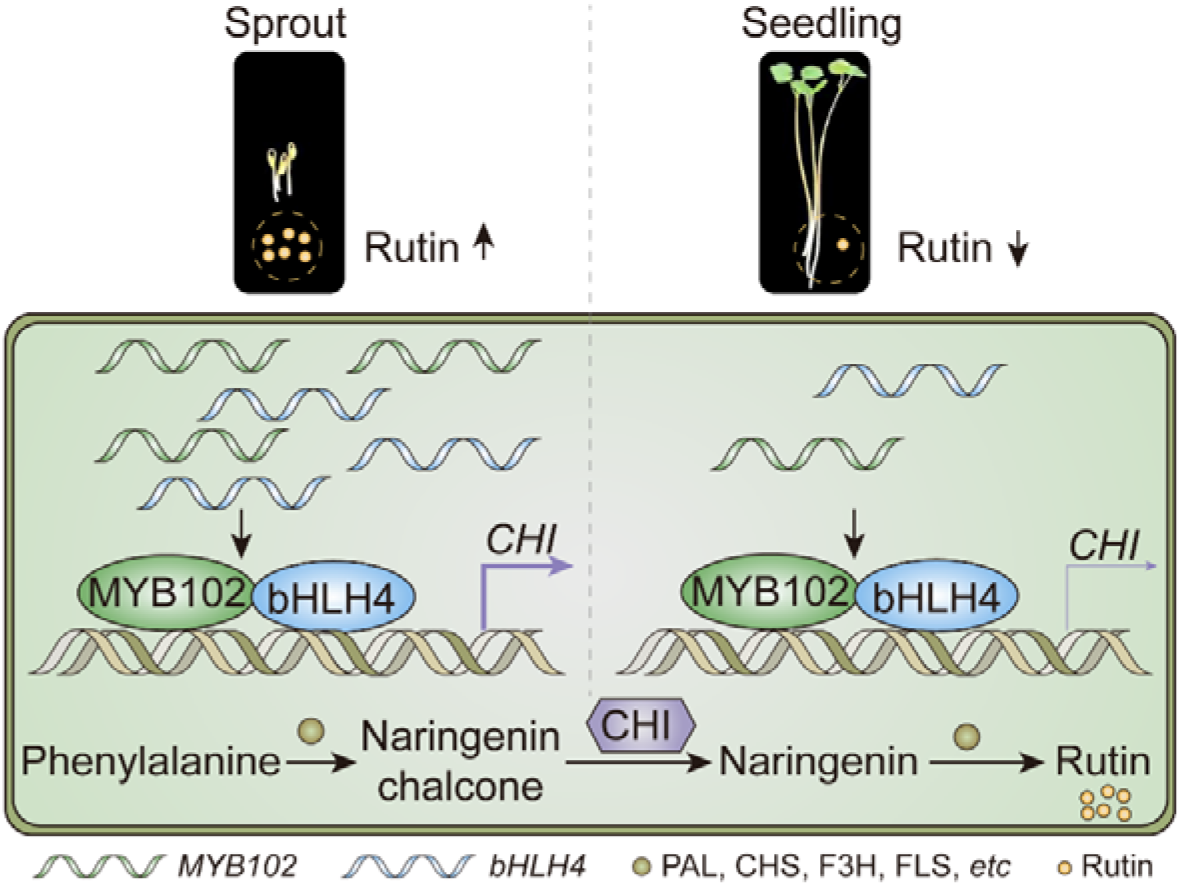
A working model of the MYB102 and bHLH4 form a transcriptional complex by inducing *CHI* expression to promote the accumulation of rutin. (Left) In sprouts, the transcripts of MYB102 and bHLH4 are abundant, thus more MYB102 and bHLH4 proteins form a complex to activate the high expression of *CHI*, since *CHI* is an important structural gene for rutin, this causes rutin levels to be high. (Right) In seedlings, unlike sprouts, MYB102 and bHLH4 transcripts are reduced, and *CHI* expression is downregulated as the number of MYB102 and bHLH4 complexes is less; thus, rutin levels are reduced.

## Acknowledgments

This work was supported by the following grants and projects: National Key R&D Program of China (2019YFC1711100, 2021YFE1011900), Scientific and Technological Innovation Project of China Academy of Chinese Medical Sciences (CI2021A03710, CI2021A041013), Opening project of Shanghai Key Laboratory of Plant Functional Genomics and Resources, National Natural Science Foundation of China (31860408, 31900258).

## References

Arce-Rodríguez M.L, Ochoa-Alejo N. 2017. An R2R3-MYB transcription factor regulates capsaicinoid biosynthesis. Plant Physiol, 174, 1359–1370.

Ariani A, Gepts P. 2015. Genome - wide identification and characterization of aquaporin gene family in common bean (Phaseolus vulgaris L.). Mol Genet Genomics, 290, 1771–1785.

Bai YC, Li CL, Zhang JW, Li SJ, Luo XP, Yao HP, Chen H, Zhao HX, Park SU, Wu Q. 2014. Characterization of two Tartary buckwheat R2R3-MYB transcription factors and their regulation of proanthocyanidin biosynthesis. Physiol Plant, 152, 431–40.

Barbehenn RV, Constabel CP. 2011. Tannins in plant–herbivore interactions. Phytochemistry, 72, 1551–1565.

Cheng AX, Han XJ, Wu YF, Lou HX. 2014. The function and catalysis of 2-oxoglutarate-dependent oxygenases involved in plant flavonoid biosynthesis. Int J Mol Sci, 15, 1080–1095.

Czechowski T, Larson TR, Catania TM, Harvey D, Brown GD, Graham IA. 2016. Artemisia annua mutant impaired in artemisinin synthesis demonstrates importance of nonenzymatic conversion in terpenoid metabolism. Proc Natl Acad Sci U S A, 113, 15150–15155.

Czemmel S, Stracke R, Weisshaar B, Cordon N, Harris NN, Walker AR, Robinson SP, Bogs J. 2009. The grapevine R2R3-MYB transcription factor VvMYBF1 regulates flavonol synthesis in developing grape berries. Plant Physiol, 151, 1513–1530.

Deluc L, Bogs J, Walker AR, Ferrier T, Decendit A, Merillon JM, Robinson SP, Barrieu F. 2008. The transcription factor VvMYB5b contributes to the regulation of anthocyanin and proanthocyanidin biosynthesis in developing grape berries. Plant Physiol, 147, 2041–2053.

Dixon RA, Lamb CJ. 1979. Stimulation of de novo synthesis of L-phenylalanine ammonia-lyase in relation to phytoalexin accumulation in Colletotrichum lindemuthianum elicitor-treated cell suspension cultures of French bean (Phaseolus vulgaris). Biochim Biophys Acta, 586, 453–463.

Dong Q, Zhao H, Huang Y, Chen Y, Wan M, Zeng Z, Yao P, Li C, Wang X, Chen H, Wu Q. 2020. FtMYB18 acts as a negative regulator of anthocyanin/proanthocyanidin biosynthesis in Tartary buckwheat. Plant Mol Biol, 104, 309–325.

Du H, Huang Y, Tang Y. 2010. Genetic and metabolic engineering of isoflavonoid biosynthesis. Appl Microbiol Biotechnol, 86, 1293–1312.

Dubos C, Stracke R, Grotewold E, Weisshaar B, Martin C, Lepiniec L. 2010. MYB transcription factors in Arabidopsis. Trends Plant Sci, 15, 573–581.

Emiliani J, Grotewold E, Ferreyra MLF, Casati P. 2013. Flavonols protect Arabidopsis plants against UV-B deleterious effects. Mol Plant, 6, 1376–1379.

Feller A, Machemer K, Braun EL, Grotewold E. 2011. Evolutionary and comparative analysis of MYB and bHLH plant transcription factors. Plant J, 66, 94–116.

Feyissa DN, Løvdal T, Olsen KM, Slimestad R, Lillo C. 2009. The endogenous GL3, but not EGL3, gene is necessary for anthocyanin accumulation as induced by nitrogen depletion in Arabidopsis rosette stage leaves. Planta 230, 747–754.

Goff SA, Cone KC, Chandler VL. 1992. Functional analysis of the transcriptional activator encoded by the maize B gene: evidence for a direct functional interaction between two classes of regulatory proteins. Genes Dev, 6, 864–875.

Gonzalez A, Zhao M, Leavitt JM, Lloyd AM. 2008. Regulation of the anthocyanin biosynthetic pathway by the TTG1/bHLH/Myb transcriptional complex in Arabidopsis seedlings. Plant J, 53, 814–827.

Hagmann ML, Heller W, Grisebach H. 1983. Induction and characterization of a microsomal flavonoid 3′-hydroxylase from parsley cell cultures. Eur J Biochem, 134, 547–554.

Hao Y, Oh E, Choi G, Liang Z, Wang ZY. 2012. Interactions between HLH and bHLH factors modulate light-regulated plant development. Mol Plant, 5, 688–697.

Holton TA, Brugliera F, Tanaka Y. 1993. Cloning and expression of flavonol synthase from Petunia hybrida. Plant J, 4, 1003–1010.

Hou S, Du W, Hao Y, Han Y, Li H, Liu L, Zhang K, Zhou M, Sun Z. 2021. Elucidation of the regulatory network of flavonoid biosynthesis by profiling the metabolome and transcriptome in Tartary Buckwheat. JAgric Food Chem, 69, 7218–7229.

Huang Y, Wu Q, Wang S, Shi J, Dong Q, Yao P, Shi G, Xu S, Deng R, Li C, Chen H, Zhao H. 2019. FtMYB8 from Tartary buckwheat inhibits both anthocyanin/proanthocyanidin accumulation and marginal Trichome initiation. BMC Plant Biol, 19, 263.

Jiang P, Burczynski F, Campbell C, Pierce G, Austria JA, Briggs CJ. 2007. Rutin and flavonoid contents in three buckwheat species Fagopyrum esculentum, F. tataricum, and F. homotropicum and their protective effects against lipid peroxidation. Food Res Int, 40, 356–364.

Jin H, Cominelli E, Bailey P, Parr A, Mehrtens F, Jones J, Tonelli C, Weisshaar B, Martin C. 2000. Transcriptional repression by AtMYB4 controls production of UV-protecting sunscreens in Arabidopsis. EMBO J, 19, 6150–6161.

Koes RE, Spelt CE, Mol JN. 1989. The chalcone synthase multigene family of Petunia hybrida (V30): differential, light-regulated expression during flower development and UV light induction. Plant Mol Biol, 12, 213–225.

Kumar S, Stecher G, Tamura K. 2016. MEGA7: molecular evolutionary genetics analysis version 7.0 for bigger datasets. Mol Biol Evol, 2016, 33, 1870–1874.

Liew CF, Loh CS, Goh CJ, Lim SH. 1998. The isolation, molecular characterization and expression of dihydroflavonol 4-reductase cDNA in the orchid, Bromheadia finlaysoniana. Plant Sci, 135, 161–169.

Li J, Luan Q, Han J, Zhang C, Liu M, Ren Z. 2020. CsMYB60 directly and indirectly activates structural genes to promote the biosynthesis of flavonols and proanthocyanidins in cucumber. Hortic Res, 7, 103.

Li P, Chen B, Zhang G, Chen L, Dong Q, Wen J, Mysore KS, Zhao J. 2016. Regulation of anthocyanin and proanthocyanidin biosynthesis by M edicago truncatula b HLH transcription factor MtTT8. New Phytol, 210, 905–921.

Lotkowska ME, Tohge T, Fernie AR, Xue GP, Balazadeh S, Mueller-Roeber B. 2015. The Arabidopsis transcription factor MYB112 promotes anthocyanin formation during salinity and under high light stress. Plant Physiol, 169, 1862–1880.

Ludwig SR, Habera LF, Dellaporta SL, Wessler SR. 1989. Lc, a member of the maize R gene family responsible for tissue-specific anthocyanin production, encodes a protein similar to transcriptional activators and contains the myc-homology region. Proc Natl Acad Sci USA, 86, 7092–7096.

Luo X, Zhao H, Yao P, Li Q, Huang Y, Li C, Chen H, Wu Q. 2018. An 2R3-MYB transcription factor FtMYB15 involved in the synthesis of anthocyanin and proanthocyanidins from Tartary buckwheat. J Plant Growth Regul, 37, 76–84.

Mona CM, Christopher JL. 1987. Chalcone isomerase cDNA cloning and mRNA induction by fungal elicitor, wounding and infection. EMBO J, 6, 1527–1533.

Montefiori M, Brendolise C, Dare AP, Lin-Wang K, Davies KM, Hellens RP, Allan AC. 2015. In the Solanaceae, a hierarchy of bHLHs confer distinct target specificity to the anthocyanin regulatory complex. J Exp Bot, 66, 1427–1436.

Mortazavi A, Williams BA, McCue K, Schaeffer L, Wold B. 2008. Mapping and quantifying mammalian transcriptomes by RNA-Seq. Nat Methods, 5, 621–8.

Negahdari R, Bohlouli S, Sharifi S, Maleki Dizaj S, Rahbar Saadat Y, Khezri K, Jafari S, Ahmadian E, Gorbani Jahandizi N, Raeesi S. 2021. Therapeutic benefits of rutin and its nanoformulations. Phytother Res, 35, 1719–1738.

Nesi N, Debeaujon I, Jond C, Pelletier G, Caboche M, Lepiniec L. 2000. The TT8 gene encodes a basic helix-loop-helix domain protein required for expression of DFR and BAN genes in Arabidopsis siliques. Plant Cell, 12, 1863–1878.

Payne CT, Zhang F, Lloyd AM. 2000. GL3 encodes a bHLH protein that regulates trichome development in Arabidopsis through interaction with GL1 and TTG1. Genetics, 156, 1349–1362.

Russell DW. 1971. The metabolism of aromatic compounds in higher plants. J Biol Chem, 246, 3870–3878.

Shang Y, Venail J, Mackay S, Bailey PC, Schwinn KE, Jameson PE, Martin CR, Davies KM. 2011. The molecular basis for venation patterning of pigmentation and its effect on pollinator attraction in flowers of Antirrhinum. New Phytol, 189, 602–615.

Spelt C, Quattrocchio F, Mol JNM, Koes R. 2000. anthocyanin1 of petunia encodes a basic helix-loop-helix protein that directly activates transcription of structural anthocyanin genes. Plant Cell, 12, 1619–1632.

Teng S, Keurentjes J, Bentsink L, Koornneef M, Smeekens S. 2005. Sucrose-specific induction of anthocyanin biosynthesis in Arabidopsis requires the MYB75/PAP1 gene. Plant Physiol, 139, 1840–1852.

Toledo-Ortiz G, Huq E, Quail PH. 2003. The Arabidopsis basic/helix-loop-helix transcription factor family. Plant Cell, 15, 1749–1770.

Vimolmangkang S, Han Y, Wei G, Korban SS. 2013. An apple MYB transcription factor, MdMYB3, is involved in regulation of anthocyanin biosynthesis and flower development. BMC Plant Biol, 13, 176.

Yang CQ, Fang X, Wu XM, Mao YB, Wang LJ, Chen XY. 2012. Transcriptional regulation of plant secondary metabolism. J Integr Plant Biol, 54, 703–712.

Yang W, Su Y, Dong G, Qian G, Shi Y, Mi Y, Zhang Y, Xue J, Du W, Shi T, Chen S, Zhang Y, Chen Q, Sun W. 2020. Liquid chromatography-mass spectrometry-based metabolomics analysis of flavonoids and anthraquinones in Fagopyrum tataricum L. Gaertn. (Tartary buckwheat) seeds to trace morphological variations. Food Chem, 331, 127354.

Yin Q, Han X, Han Z, Chen Q, Shi Y, Gao H, Zhang T, Dong G, Xiong C, Song C, Sun W, Chen S. 2020. Genome-wide analyses reveals a glucosyltransferase involved in rutin and emodin glucoside biosynthesis in Tartary buckwheat. Food Chem, 318, 126478.

Wang L, Deng R, Bai Y, Wu H, Li C, Wu Q, Zhao H. 2022. Tartary Buckwheat R2R3-MYB gene FtMYB3 negatively regulates anthocyanin and proanthocyanin biosynthesis. Int. J Mol Sci. 23, 2775.

Wang N, Xu H, Jiang S, Zhang Z, Lu N, Qiu H, Qu C, Wang Y, Wu S, Chen X. 2017. MYB12 and MYB22 play essential roles in proanthocyanidin and flavonol synthesis in red - fleshed apple (Malus sieversii f. niedzwetzkyana). Plant J, 90, 276–292.

Williams CA, Grayer RJ. 2004. Anthocyanins and other flavonoids. Nat Prod Rep, 21, 539–573.

Zhang D, Jiang C, Huang C, Wen D, Lu J, Chen S, Zhang T, Shi Y, Xue J, Ma W, Xiang L, Sun W, Chen S. 2019. The light-induced transcription factor FtMYB116 promotes accumulation of rutin in Fagopyrum tataricum. Plant Cell Environ, 42, 1340–1351.

Zhang K, Logacheva MD, Meng Y, Hu J, Wan D, Li L, Janovská D, Wang Z, Georgiev MI, Yu Z, Yang F, Yan M, Zhou M. 2018. Jasmonate-responsive MYB factors spatially repress rutin biosynthesis in Fagopyrum tataricum. J Exp Bot, 69, 1955–1966.

Zhang LJ, Li XX, Ma B, Gao Q, Du HL, Han Y, Li Y, Cao Y, Qi M, Zhu Y, Lu H,Ma M, Liu L, Zhou J, Nan C, Qin Y, Wang J, Cui L, Liu H, Liang C, Qiao Z. 2017. The Tartary buckwheat genome provides insights into rutin biosynthesis and abiotic stress tolerance. Mol Plant, 10, 1224–1237.

Zhou LL, Shi MZ, Xie DY. 2012. Regulation of anthocyanin biosynthesis by nitrogen in TTG1-GL3/TT8-PAP1-programmed red cells of Arabidopsis thaliana. Planta, 236, 825–837.

Zhou M, Sun Z, Wang C, Zhang X, Tang Y, Zhu X, Shao J, Wu Y. 2015. Changing a conserved amino acid in R2R3-MYB transcription repressors results in cytoplasmic accumulation and abolishes their repressive activity in Arabidopsis. Plant J, 84, 395–403.

Zhou M, Zhang K, Sun Z, Yan M, Chen C, Zhang X, Tang Y, Wu Y. 2017. LNK1 and LNK2 corepressors interact with the MYB3 transcription factor in phenylpropanoid biosynthesis. Plant Physiol, 174, 1348–1358.

Zhou M, Memelink J. 2016. Jasmonate-responsive transcription factors regulating plant secondary metabolism. Biotechnol Adv, 34, 441–449.

Zhou M, Sun Z, Ding M, Logacheva MD, Kreft I, Wang D, Yan M, Shao J, Tang Y, Wu Y, Zhu X. 2017. FtSAD2 and FtJAZ1 regulate activity of the FtMYB11 transcription repressor of the phenylpropanoid pathway in Fagopyrum tataricum. New Phytol, 216, 814–828.

Zimmermann IM, Heim MA, Weisshaar B, Uhrig JF. 2004. Comprehensive identification of Arabidopsis thaliana MYB transcription factors interacting with R/B-like BHLH proteins. Plant J, 40, 22–34.

